# The influence of social lifestyles on host-microbe symbioses in the bees

**DOI:** 10.1101/2023.06.12.544601

**Authors:** Lauren Mee, Seth M Barribeau

## Abstract

Microbiomes are increasingly recognised as critical for the health of an organism. In eusocial insect societies, frequent social interactions allow for high fidelity transmission of microbes across generations, leading to closer host-microbe coevolution. The microbial communities of bees with different social lifestyles are less well studied, and few comparisons have been made between taxa that vary in social structure. To address this gap, we leveraged a cloud-computing resource and publicly available transcriptomic data to conduct a survey of microbial diversity in bee samples from a variety of social lifestyles and taxa. We consistently recover the core microbes of well-studied corbiculates, supporting this method’s ability to accurately characterise microbial communities. We find that the bacterial communities of bees are influenced by host location, phylogeny, and social lifestyle, although no clear effect was found for fungal or viral microbial communities. Bee genera with more complex societies tend to harbour more diverse microbes, with *Wolbachia* detected more commonly in solitary tribes. We present the first description of the microbiota of Euglossine bees and find that they do not share the “corbiculate” core microbiome. Notably, we find that bacteria with known anti-pathogenic properties are present across social bee genera, suggesting that symbioses that enhance host immunity are important with higher sociality. Our approach provides an inexpensive means of exploring microbiomes of a given taxa and identifying avenues for further research. These findings contribute to our understanding of the relationships between bees and their associated microbial communities, highlighting the importance of considering microbiome dynamics in investigations of bee health.

## 2 Introduction

In the insect world, microbial symbionts can play a major role in many biological processes [Munoz-Benavent et al., 2021], including reproduction [Bourtzis et al., 1996; Singh and Linksvayer, 2020; Werren et al., 2008], nutrition [Andersen et al., 2012; Cheng et al., 2019] and pathogen defense [Benoit et al., 2017; Bian et al., 2010; Duplouy et al., 2015]. For social insects, where consistent social contact between conspecifics allows for high-fidelity vertical transmission of microbial communities, these symbionts can be passed on for generations, allowing for coevolution of microbiome and host [Dietrich et al., 2014; Kwong et al., 2017b; Lombardo, 2008; Sanders et al., 2014; Zhang and Zheng, 2022]. This has been demonstrated in the obligately eusocial corbiculate bees, which all share a core set of bacterial microbes [Koch and Schmid-Hempel, 2011; Koch et al., 2013; Kwong and Moran, 2016; Kwong et al., 2017b; Lim et al., 2015; Moran et al., 2012]. The members of this conserved bacterial complement are important for the health of their hosts, particularly by protecting against infectious disease [Anderson et al., 2014; Bonilla-Rosso and Engel, 2018; Koch and Schmid-Hempel, 2012; Miller et al., 2021; Véasquez et al., 2012].

However, there are very few bee microbial studies outside of these eusocial corbiculates [Handy et al., 2022; Kapheim et al., 2021; McFrederick et al., 2012, 2014, 2017], meaning the microbiomes of most less studied bee species remain a mystery. One of the current approaches of characterising the microbiome of a host is to use metagenomic Next Generation Sequencing (mNGS), where all DNA (or RNA) from a given environment – i.e. an insect gut – is sequenced and the microbial community characterised. While the cost of producing NGS data has dramatically reduced over recent years, it remains reasonably expensive, taking into account sample extraction, library production, sequencing costs, and having the appropriate informatics infrastructure in order to store, process and analyse data [Krampis and Wultsch, 2015].

One attractive solution for some analyses is to use cloud-computing resources [Krampis and Wultsch, 2015]. CZID.org, for example, is an approachable, open source cloud-based service which can provide microbial identification for many different sample types and host species [Kalantar et al., 2020]. Here we use this approach to examine NGS datasets from 18 bee genera spanning 100 million years of divergence (Figure 1, Peters et al. [2017]) that vary in their social structure, ranging from solitary to obligately eusocial. As one of the purposes of this analysis was to assess any differences in microbial composition potentially caused by consistent social interaction, we simplified the many different distinctions in social structure found in the literature to: 1) solitary, where species do not provide any brood care and associate with conspecifics only for mating; 2) facultatively eusocial, which included any species that had considerable contact with conspecifics (i.e. communal nesting) and some brood care (primitively eusocial) but where individuals can and do live solitarily and 3) obligately eusocial species that only ever exhibit eusocial behaviours and solitary living is impossible (Figure 1). We used this framework to systematically test whether social structure, location or bee taxa affect microbial composition across the bees.

**Figure 1:**
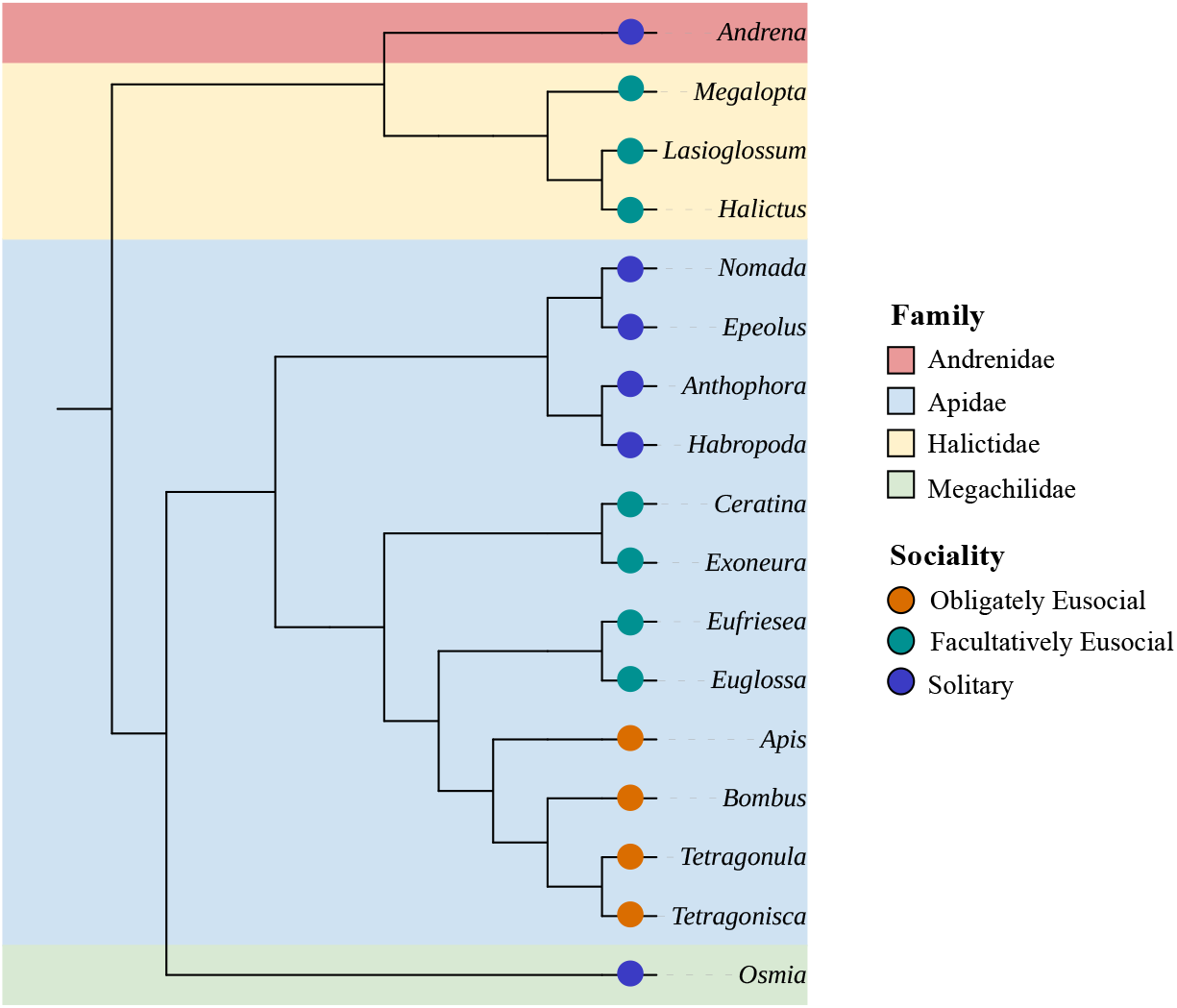
Phylogeny of the genera included in these analyses coloured by family, with the sociality of each genus specified by a coloured circle. The corbiculate bees are marked within a black lined box. This tree is based on accepted topology in the literature [Bossert et al., 2019; da Silva, Jack, 2021; Gibbs et al., 2012; Husemann et al., 2021; Kapheim et al., 2019; Lu et al., 2021]. Branch lengths are not indicative of evolutionary time. Figure produced using iTol’s web tool [Letunic and Bork, 2021].

## 3 Materials and methods

### 3.1 Sample selection

We analysed sequence data sourced from NCBI’s Sequence Reads Archive (SRA, Katz et al. [2022]; Kodama et al. [2012]; Leinonen et al. [2010]), accessed September 2022. We included all available RNA-Seq adult bee samples that included the animal’s abdomen (including pooled individuals) and we excluded projects that exclusively sequenced any other part (e.g. antennae, brain, ovaries). We only included unaltered control specimens (i.e. no treatment or stressor introduced/administered) to ensure that the microbial composition was as natural as possible.

### 3.2 Processing, mapping and uploading reads

All sequence data (fastq format, Table 1, see **S4.1**) were downloaded and unpacked from the SRA using prefetch and fasterq-dump from the SRA-toolkit (version 3.0.0, Katz et al. [2022]; Kodama et al. [2012]; Leinonen et al. [2010]). From here we split the pipeline: files sequencing the European honey-bee *Apis mellifera* were uploaded directly to CZID.org using the command-line interface (version 4.1.2), and non-*A. mellifera* sequences were retained for further processing. CZID (Chan Zuckerberg ID, previously known as IDSeq, Kalantar et al. [2020]) is a cloud-based, open-source platform that maps input sequence files against a chosen species genome and then aligns any unmapped reads to NCBI databases in order to detect non-host sequences (see reference for pipeline details). In each of these non-host taxa “hits” the number of reads are recorded and these counts can be considered as representative of microbial taxa presence and abundance. The genome that sequences will be mapped against is selected from a pre-determined list and at the time of the analysis (October 2022) the host genome option “Bee” included only the honeybee, *A. mellifera*, genome. Therefore, non-*A. mellifera* samples required a number of pre-processing steps.

First, each sample was assigned the phylogenetically closest reference genome (see **S1**). These sequence files were then mapped against each respective genome using STAR (version 2.7.10a, Dobin and Gingeras [2016]; Dobin et al. [2013]). Every sample that achieved *>* 50% of reads successfully mapping to the reference genome proceeded to the next step. For the samples that had *≤* 50% reads fail to map because they were ‘too short’, we repeated the map-ping with slightly relaxed parameters (--outFilterScoreMinOverLread 0.3 --outFilterMatchNminOverLread 0.3). This was needed when the species was comparatively phylogenetically distant from the nearest available genome. Regardless of the success of the second mapping run, all unmapped sequence files were then uploaded to CZID.org for taxonomic assignment using pipeline version 7.1.

**Table 1:**
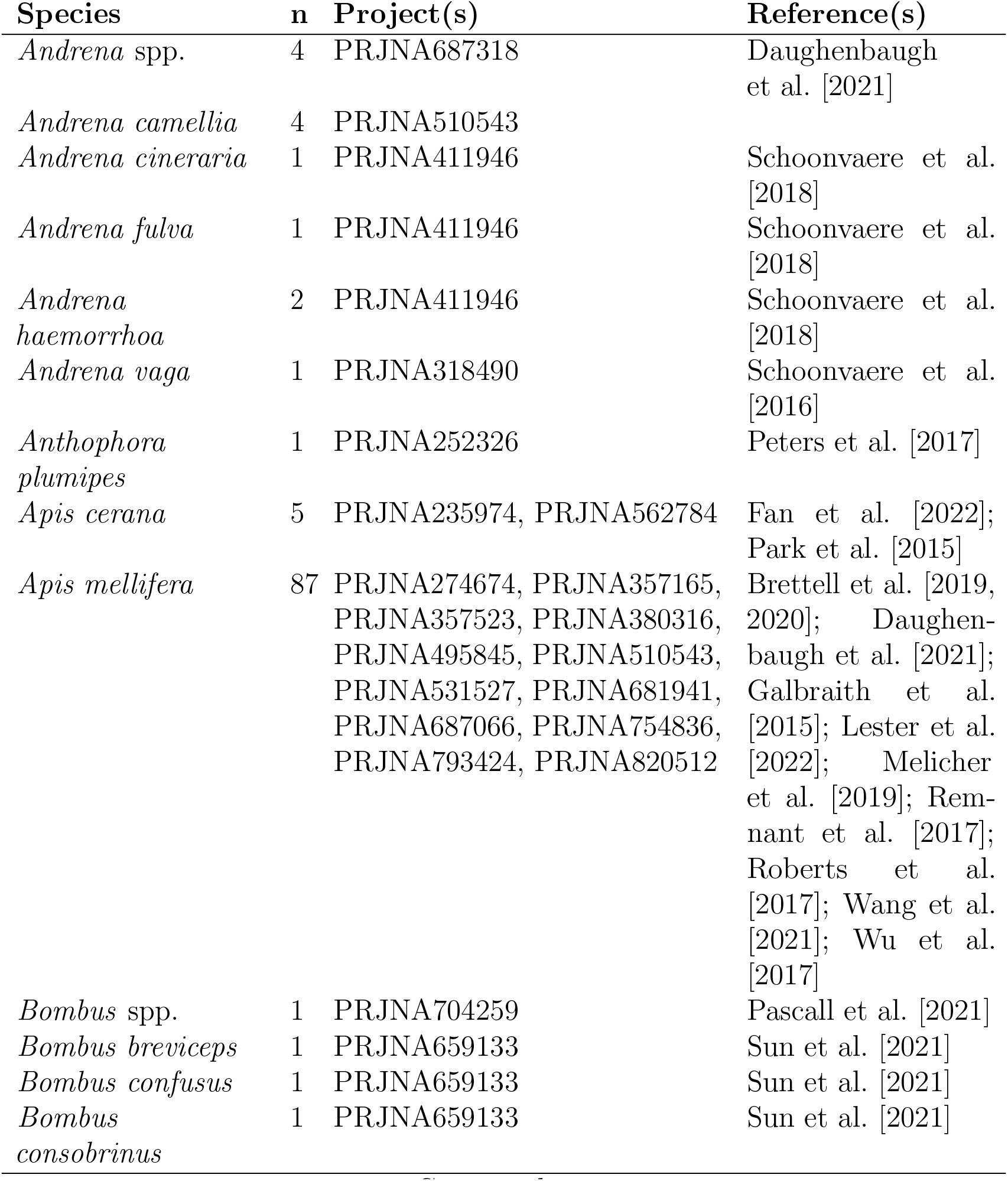
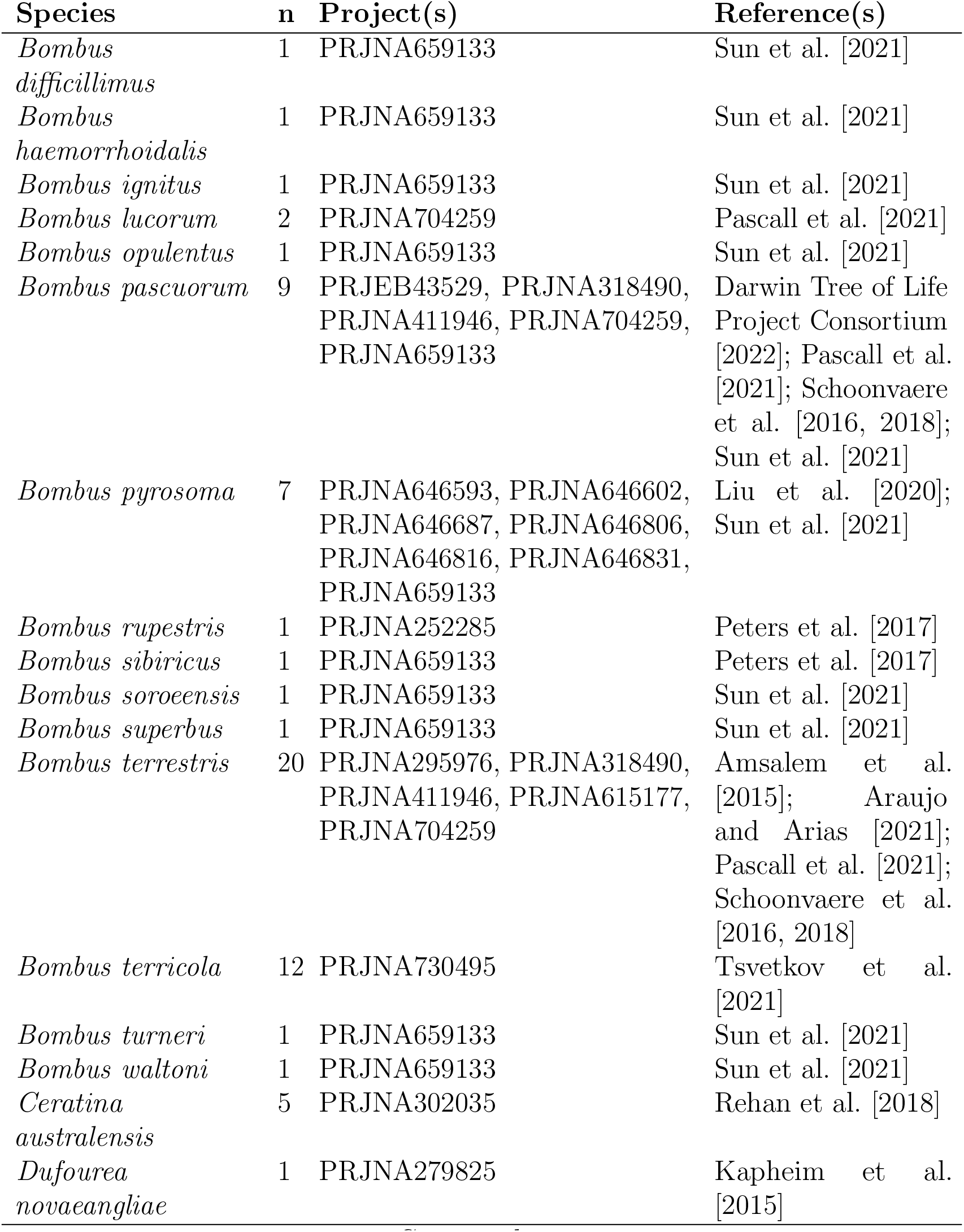
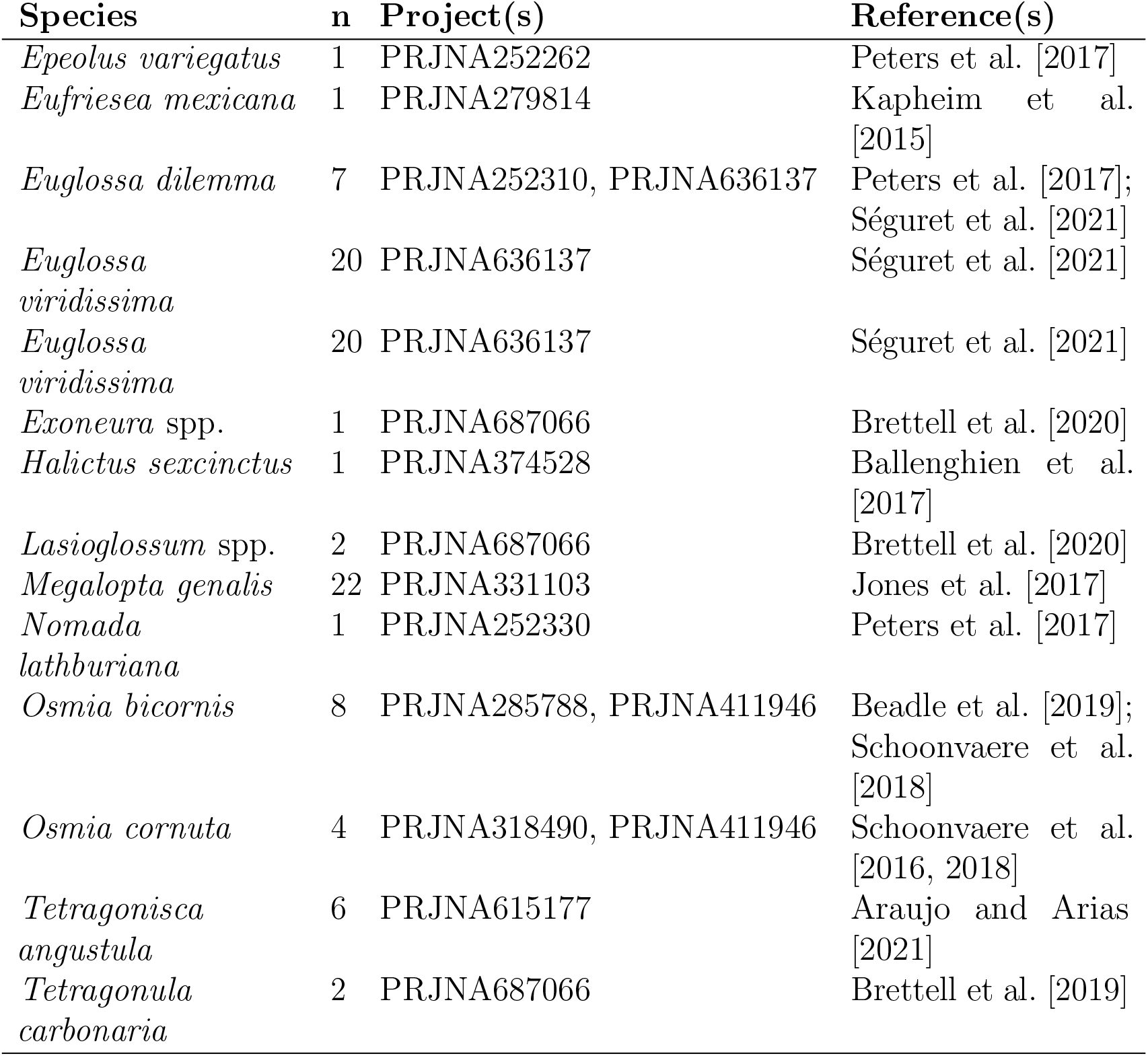
List of host species with associated NCBI projects and references when available. See **S2** for further details.

### 3.3 Taxonomy

All taxonomic classifications of the identified microbes were sourced from the NCBI Taxonomy (taxonomy dump file from NCBI ftp service Federhen [2012]; Schoch et al. [2020], accessed 18th October 2022). A single manual change was made: we distinguish the *Lactobacillus Firm-5* as a separate genus to *Lactobacillus*, as this taxonomic cluster has repeatedly been found to be an important member of the corbiculate bee microbiome [Kwong et al., 2017b; Martinson et al., 2011; Véasquez et al., 2012].

CZID also uses the NCBI taxonomy as the basis of its taxon reports, but, as it is only updated periodically, there were some minor differences between taxa identified as hits by CZID and corresponding classifications in the NCBI taxonomy dump file. In these instances, we updated the taxon reports to reflect the more recent classifications (NCBI). For all analyses, we only used genus-level CZID results (i.e. the evalue, aggregate score, read count and read reads per million [rPM]) as species information was not available for all taxa. To collapse species to the genus level we took the minimum, maximum and sums of the evalue, aggregate score and read counts/rPM, for all species within a genus. To control for potential contamination, CZID uses a “blank” as background to compute a taxon level z-score which reflects the likelihood of a taxonomic hit being a contaminant. As these experiments are from many different laboratories using different reagent kits throughout extraction and sequencing, we selected a generic water as the blank sample as it is likely to be analogous to other molecular grade waters used in sample preparation (specifically, “EARLI Novaseq Water Control”).

### 3.4 Generating community count tables

Each CZID taxon report file is produced individually per host sample. Each report file was checked for taxa that matched to non-microbial sources – such as the host, other invertebrates or plants – and removed when found. These files were then iterated through and non-host taxon hits were filtered according to the following criteria: 1) read counts were present above 5 reads per million, 2) alignment length was larger than 50 nucleotides, 3) e-value was below 1*e −* 6, 4) CZID aggregate and z-scores were above 0, and 5) alignment percent identity was above 90%. This process was run separately for bacteria, eukaryote, and viral taxa hit sequences. CZID aligns suspected non-host reads to both the NCBI nucleotide (NT) and non-redundant protein (NR) sequence databases. For prokaryotic and eukaryotic taxa, the above filters were assigned to the taxa hits mapped against the NT database; the viral taxa were assessed against the NR database results. This is necessary as viruses evolve so rapidly that they can fail to map to the NT database but map perfectly well against the more conserved NR database. Viral taxa were analysed at family level, with bacterial and eukaryote taxa at genus level. Results of each host sample were combined into a single counts table per microbial classification (bacteria, eukaryotes, and viruses).

### 3.5 Beta diversity (dissimilarity) analyses

Count tables were further reduced by removing host samples that had fewer than 100 non-host reads total and microbial taxa that were present in less than 5% of the remaining samples. As sample phylogeny was to be considered in microbial composition, we restricted sample sets to taxa that contained at least four samples to allow for centroid calculation. Host taxa with fewer samples were removed. In the bacterial analysis, this could be done to the level of host tribe, and in the other two analyses, host family.

Beta diversity was calculated with vegan (version 2.6-4, Dixon [2003]) in R (version 4.2.2, R Core Team [2020]) and its associated functions. Bray-Curtis dissimilarity matrices were calculated for each microbial category using the function avgdist with 10,000 iterations. Rarefaction for each matrix was set to use the lowest number of reads from the smallest sample grouping of sociality – solitary – in order to retain as many samples of that grouping as possible. This read limit was therefore different for each of the three matrices: bacteria *n* = 323, eukaryotes *n* = 171, viruses *n* = 111. Samples with total reads less than this number were discarded. For the virus analysis, two further samples were removed to ensure there were no singletons within social lifestyle, continent or host family factor levels. Rarefied reads were used to make 10,000 distance matrices and the final matrix consisted of the average distances computed across these iterations.

Non-metric multidimensional scaling (NMDS) was used to visualise dissimilarities, computed by metaMDS. To assess whether variables of interest – social lifestyle, phylogeny, location – significantly affected community composition we performed permutational multivariate analyses (PERMANOVA) using adonis2 with 9,999 permutations. Each factor was checked for homogeneity of group dispersion using betadisper to compute average distances around the median and ANOVA to test significance of any difference between groups.

### 3.6 Predicting microbial complements

We assessed filtered count data for each microbial grouping to determine prevalence of microbial taxa per host species. Bacterial data was subject to further scrutiny where each tribe of bees was assessed for average relative abundance and prevalence of all detected prokaryotic species. Those at above 50% prevalence and 0.01% average relative abundance per tribe were considered potential members of conserved tribe-level community, termed here as an “associate” species. Overlaps of prokaryotic species by sample tribe, family and sociality was also considered. Finally, hosts were checked specifically to see if they contained any of the core phylotypes found associated with corbiculate bees in previous studies. Prevalence was calculated per tribe for the corbiculates (Apini, Bombini, Meliponini and Euglossini), with non-corbiculates ordered by sociality.

## 4 Results

### 4.1 Sample selection and CZID pipeline

There were initially 285 bee samples that met the selection requirements for download from the SRA. After filtering out samples that had too few counts after host mapping (in non-*A. mellifera* samples), the CZID pipeline, and further filtering steps, there were 254 samples remaining, containing bee tissue from 4 phylogenetic families (Figure 1), 14 tribes, 18 genera and 45 species from experiments across six continents (Table 1, see **S2**). There were considerably more *Apis* and *Bombus* samples available and included (92 and 65 samples respectively) and 79.9% of all samples were from the Apidae family, particularly from corbiculate species. 165 samples are obligately eusocial, 59 facultatively eusocial, and 30 solitary. All samples successfully ran through the CZID pipeline (version 7.1), with 97% passing quality control with more than 50% of input reads (see **S3**).

### 4.2 Detected microbial community

#### 4.2.1 Bacteria

There were sufficient reads in 227 samples from 10 bee genera resulting in the detection of 65 prokaryotic taxa (Figure 2, see **S4**). The most taxa-rich host family was Apidae, which had unique taxa, while all taxa detected in other families were also present in Apidae (see **Supplementary Figure 1**). There were no bacterial taxa found only in solitary hosts, whereas there were 1 and 11 taxa unique to facultatively and obligately eusocial hosts, respectively. The former was *Asticcacaulis*, an associate bacterial taxa of Euglossini samples (Table 2), and the latter consisted of *Lactobacillus: Firm-5*, *Bartonella*, *Apibacter*, *Alcaligenes*, *Brevibacterium*, *Citrobacter*, *Deinoccocus*, *Enterobacter*, *Orbus*, *Prevotella* and *Shigella*. The majority of detected taxa belong to the Proteobacteria phylum.

**Figure 2:**
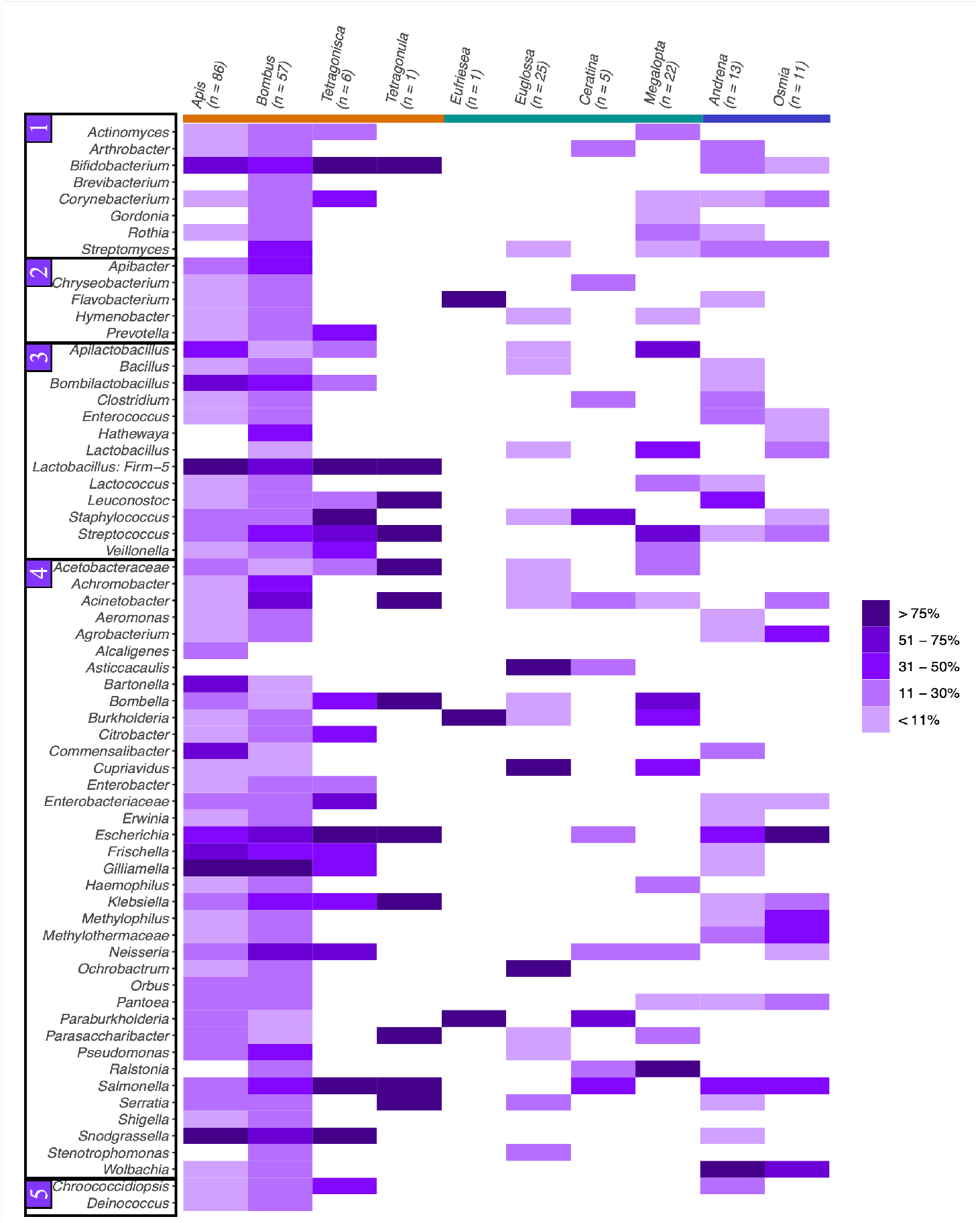
Heatmap of bacterial prevalence in each genus of host. Bacterial taxa are ordered 1) Actinobacteria, 2) Bacteroidota, 3) Firmicutes, 4) Proteobacteria and 5) other. Host genera are coloured by sociality: orange = obligately eusocial, green = facultatively eusocial, blue = solitary.

#### 4.2.2 Eukaryotic and viral taxa

There were considerably fewer samples available for determining eukaryote and viral composition after filtering steps (see **S5-6**). In 158 samples, we identified 32 eukaryotic taxa, including 24 fungi and five genera from the parasitic family Trypanosomatidae (see **Supplementary Figure 2**). The two fungal genera *Alternaria* and *Aspergillus* were detected in the majority of species, appearing in 13 and 11 out of 17 species, respectively. 12 viral families – six of which from the phylum Pisuviricota – were found across 88 host samples (see **Supplementary Figure 3**).

### 4.3 Differences in microbial composition

For the viruses, there was no significant effect of sociality, host family or continent where the sample was collected on the data (Figure 3:A,D,G, see **S7**). In eukaryotes (Figure 3:C,F,I), sociality and continent were statistically significant factors (sociality: pseudo-*F* = 2.271, *p* = 0.001; continent: pseudo-*F* = 1.794, *p* = 0.001), but both are overdispersed, suggesting caution in interpreting these results (see **S7**).

**Figure 3:**
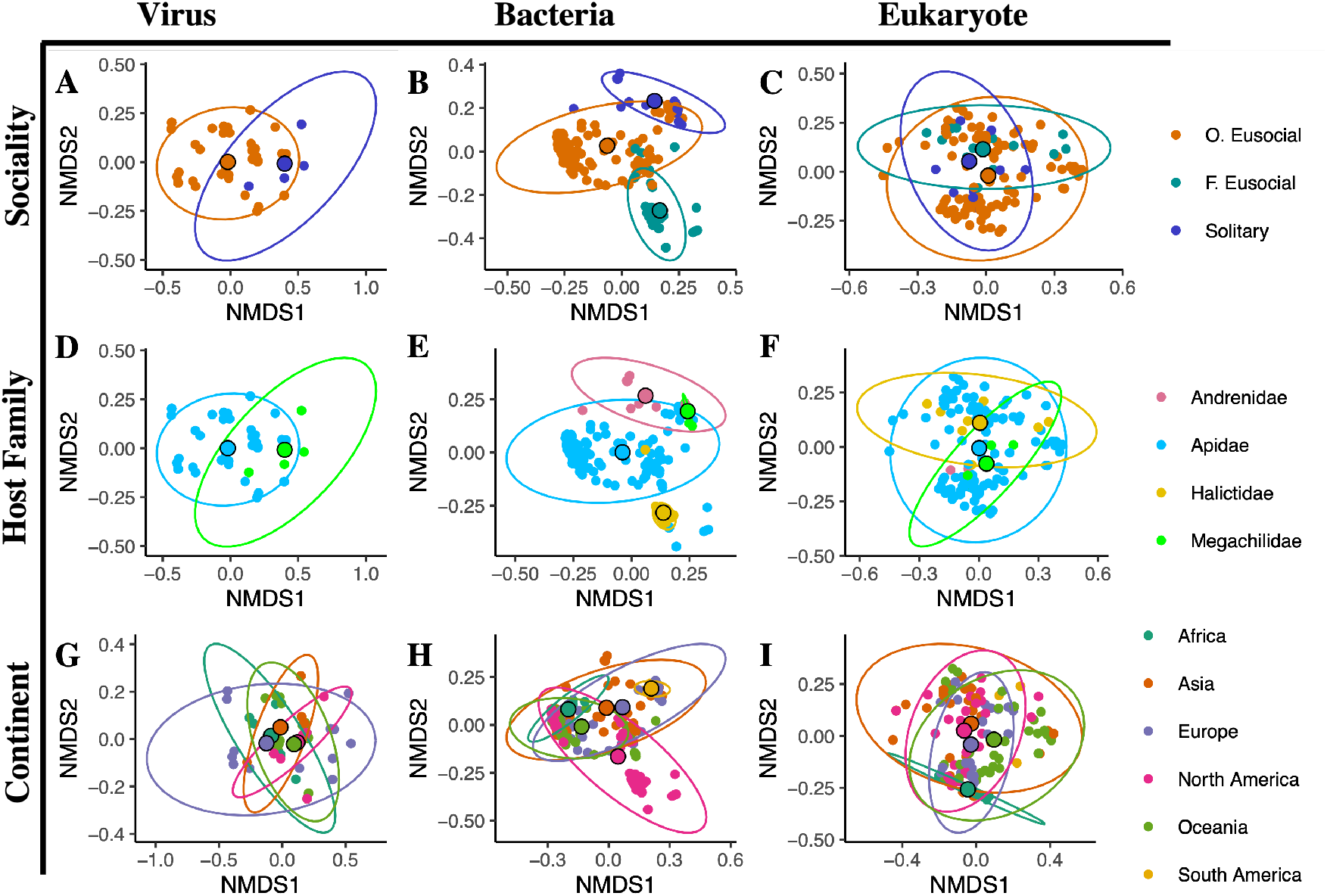
NMDS plots of Bray-Curtis dissimilarity matrices computed separately for virus (column 1), bacterial (column 2), and eukaryote (column 3) reads. Three factors were tested to assess influence on composition: sociality (row 1), host family (row 2) and continent where the samples were collected according to NCBI SRA records (row 3). Centroids for each factor level are shown larger and bordered in black. Axes may differ to incorporate full ellipses.

Sociality significantly influences bacterial composition (Figure 3:B), has homogeneous dispersion (see **S7**) and significantly influences the composition of the distance matrix (pseudo-*F* = 2.884, *p* = 0.001). This was mostly driven by the differences between obligately and facultatively eusocial samples (Pairwise PERMANOVA: *p* = 0.0195, Benjamini-Hochburg correction, see **S8**). Host family and continent (Figure 3:E,H) both also significantly affected bacterial composition (pseudo-*F* = 4.318, *p* = 1*e −* 04 and pseudo-*F* = 2.361, *p* = 1*e −* 04, respectively), and are unaffected by heterogeneous dispersion (see **S8** for pairwise PERMANOVA).

#### 4.3.1 Tribe-bacterial associates

In the more commonly studied corbiculate tribes – Apini, Bombini and Meliponini – we find at least two previously described “corbiculate” core phylotypes as associate taxa (Table 2). All eight of the taxa associated with Apini are included in the core phylotypes. All three of these tribes share an association with *Snodgrassella*, yet there is no overlap between associate taxa of these three and the other corbiculate tribe, Euglossini. *Wolbachia* is an associate of the two solitary tribes included in this analysis.

**Table 2:** Associate bacterial taxa found at above 50% prevalence and 0.01% relative abundance per tribe. Tribe cells are coloured according to host family: blue for Apidae, red for Andrenidae, yellow for Halictidae, green for Megachilidae. “Corbiculate” core bacterial taxa are indicated with *^∗^*. Only tribes included in the bacterial dissimilarity matrix were assessed (see **S1).**

| Tribe | n | Associate Taxa |
| --- | --- | --- |
| Andrenini | 13 | *Wolbachia* |
| Apini | 86 | *Bartonella* *, *Bifidobacterium* *, *Bombilactobacillus* *, *Commensalibacter* *, *Frischella* *, *Gilliamella* *, *Lactobacillus: Firm-5* *, *Snodgrassella* * |
| Augochlorini | 22 | *Apilactobacillus* *, *Bombella* *, *Ralstonia*, *Streptococcus* |
| Bombini | 57 | *Acinetobacter*, *Escherichia*, *Gilliamella* *, *Lactobacillus: Firm-5* *, *Snodgrassella* * |
| Ceratini | 5 | *Paraburkholderia*, *Staphylococcus* |
| Euglossini | 26 | *Asticcacaulis*, *Cupriavidus*, *Ochrobactrum* |
| Meliponini | 7 | *Bifidobacterium* *, *Escherichia*, *Snodgrassella* *, *Staphylococcus* |
| Osmiini | 11 | *Escherichia*, *Wolbachia* |

#### 4.3.2 Corbiculate core taxa

We find that the “corbiculate” core bacterial taxa are widely prevalent in the three well studied eusocial tribes: Apini, Bombini and Meliponini (Figure 4, see **S9**). This pattern was not repeated in Euglossini, however, where only *Apilactobacillus* was detected at low average relative abundance and prevalence. *Apilactobacillus* was interestingly found at considerable prevalence in facultatively eusocial hosts, specifically in the *Megalopta* genus, where it was detected in 15/22 samples. Other bacterial phylotypes were detected in three solitary samples: *Bifidobacterium* was detected in one individual *Andrena haemorrhoa* sample (SRR6148367), an individual *Osmia cornuta* (SRR6148371) and in a sample of pooled *Andrena* individuals of different species (SRR13404633). In the latter, *Gilliamella*, *Snodgrassela*, *Bombilacto-bacillus* and *Frischella* were also detected. *Bombiscardovia* and *Candidatus Schmidhempelia*, both taxa previously found associated with *Bombus* bees, were not detected in the analysis after filtering. *Apilactobacillus*, *Bombella* and *Parasaccharibacter* were at considerable prevalence in the facultatively eusocial bees. These values are driven largely by *Megalopta* samples (Figure 2).

**Figure 4:**
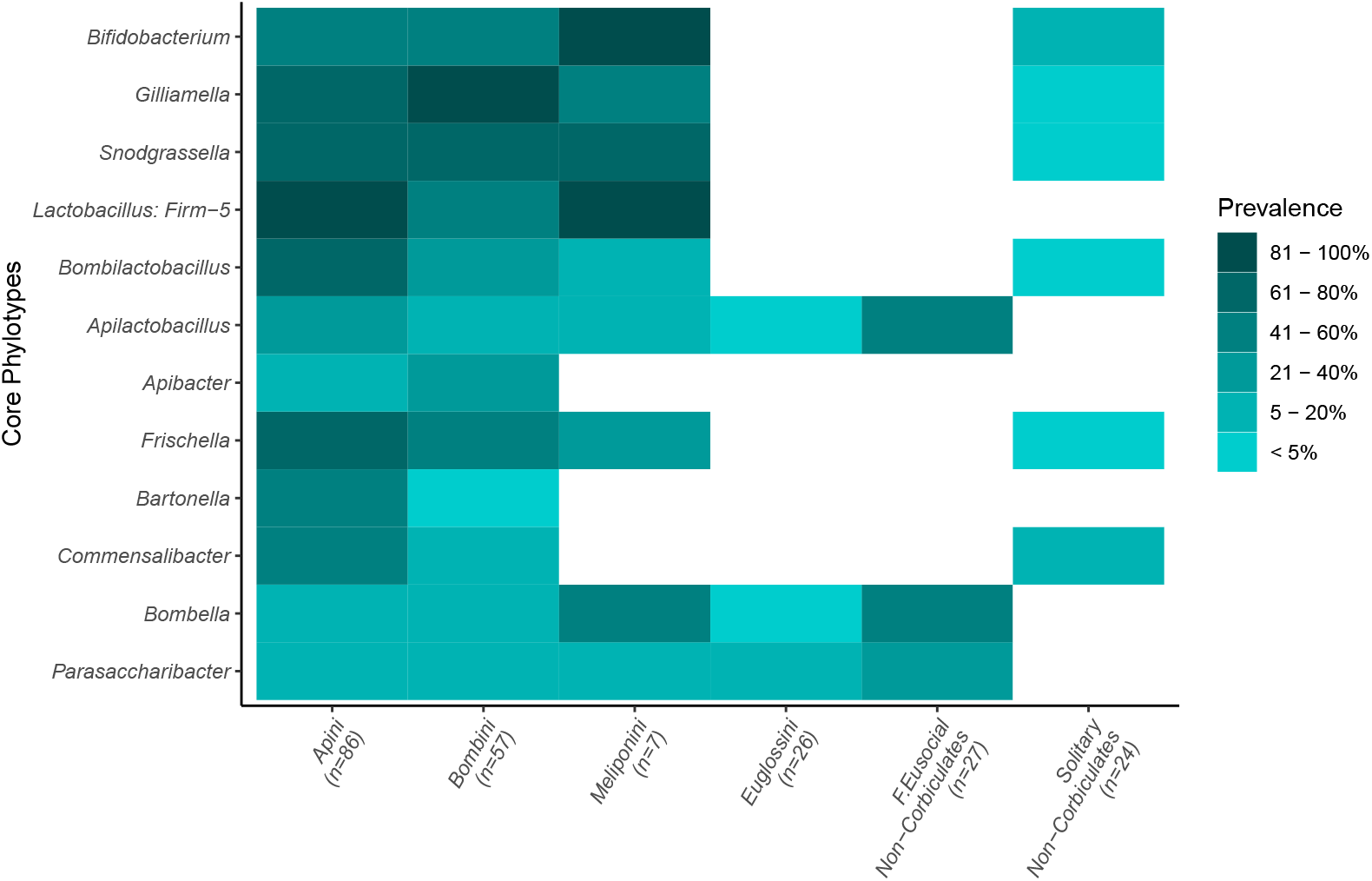
Prevalence of different microbial taxa previously described in the literature as part of the “corbiculate” core bacteria across samples. Darker tiles indicate higher prevalence.

## 5 Discussion

### 5.1 Bacterial complement affected by location, phylogeny and sociality

We find that bacterial communities are significantly affected by social lifestyle, family and collection location of the bee (Figure 3:B,E,H, see **S7**). Location and phylogeny has been found to be significant drivers of bee bacterial communities elsewhere, but there isn’t always consensus on which is more important. While some studies can identify communities to specific subfamilies or even species [Dew et al., 2020; Kwong and Moran, 2015; Kwong et al., 2017b], others find location to be more informative [Kapheim et al., 2021; Keller et al., 2013; McFrederick and Rehan, 2016, 2019; McFrederick et al., 2017], though often both play a significant role [McFrederick et al., 2012; Shell and Rehan, 2022].

It is likely that the contribution of each factor is further determined by the social lifestyle of the bee: social living allows for transmission of symbiont species in eusocial insect societies, where this vertical transmission route allows for coevolution of unique and long-lasting host-microbe associations [Dietrich et al., 2014; Kwong et al., 2017b; Lombardo, 2008; Sanders et al., 2014; Zhang and Zheng, 2022]. Solitary animals, on the other hand, are likely to have less stable communities that are largely acquired from the immediate environment [Voulgari-Kokota et al., 2019]. We see this in some of the obligately eusocial samples: when the bacterial community NMDS plots are clustered by tribe, there is a clear group of Apini samples to the left of the NMDS1 (Figure 5), despite the fact that these samples came from 11 different countries across five continents (Table 1, see **S1**).

**Figure 5:**
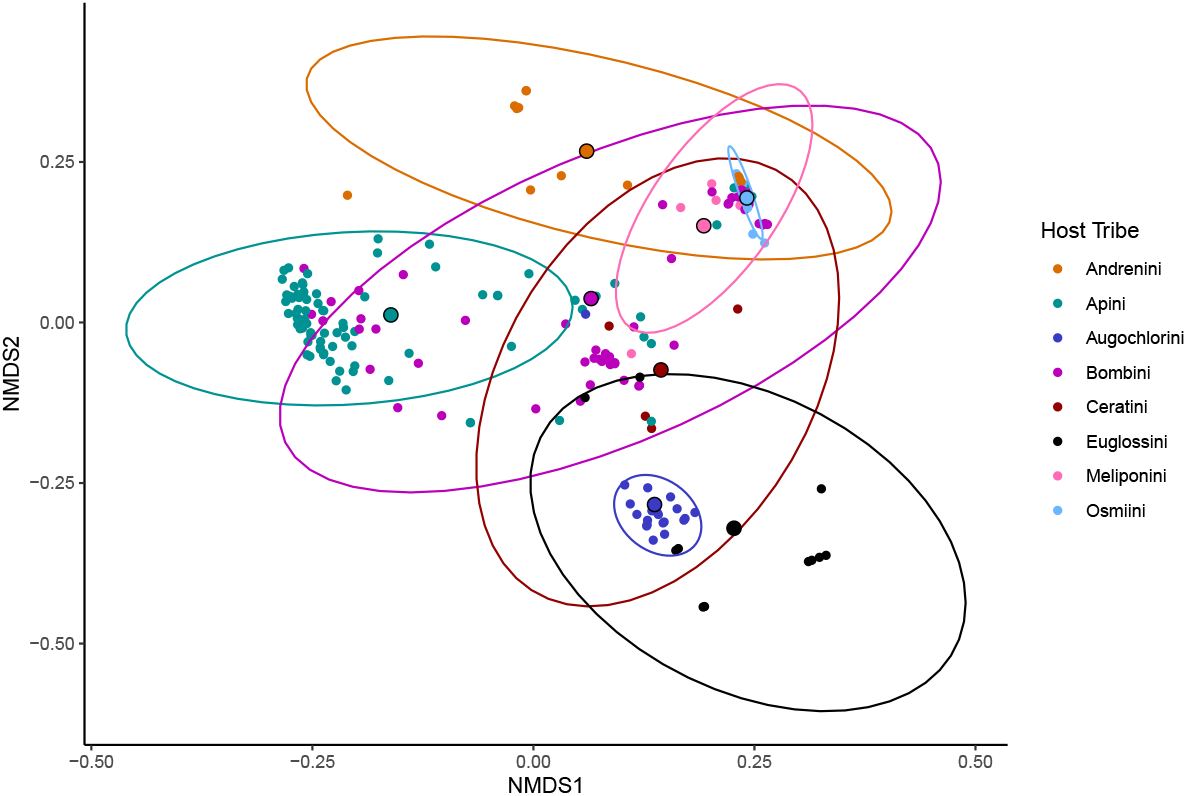
NMDS plot of Bray-Curtis dissimilarity matrix computed for bacterial read counts shows clustering of samples by host tribe. Centroids for each factor level are shown larger and bordered in black.

The limited availability of samples from solitary species does, in turn, somewhat limit the ability to untangle the microbial community composition of solitary bees. Specifically, the samples we analysed from solitary tribes Andrenini and Osmiini were primarily derived from a handful of studies (see **S1**). However, if solitary species have microbiomes that are predominantly environmentally acquired and lack the consistent vertical transmission of eusocial bees, then they should be more variable and show greater dispersion around the median than more social groups. We do find this (see **S7**), but the differences in variance is small and non-significant. Future work that includes more solitary samples would be able to better test whether solitary species have more variable microbial communities than the well characterised and more strongly vertically transmitted social microbiomes.

### 5.2 Social lifestyle impacts number and type of associate taxa

Tribes made of obligately eusocial species have the most associate microbe species in this analysis (Table 2, Figure 6:A), with Apini, Bombini and Meliponini being associated with eight, five and four bacterial genera respectively. Of these, at least two bacterial taxa were from the identified “corbiculate” core per tribe (Figure 4). This again lends weight to the hypothesis that vertical transmission leads to more stable communities in the social bees, allowing for the establishment of multiple fixed associations. We also detect more associated bacterial genera with increasing number of samples (Figure 6:B), though it should be mentioned that Meliponini has double the identified associate taxa from fairly few samples relative to the solitary tribes.

**Figure 6:**
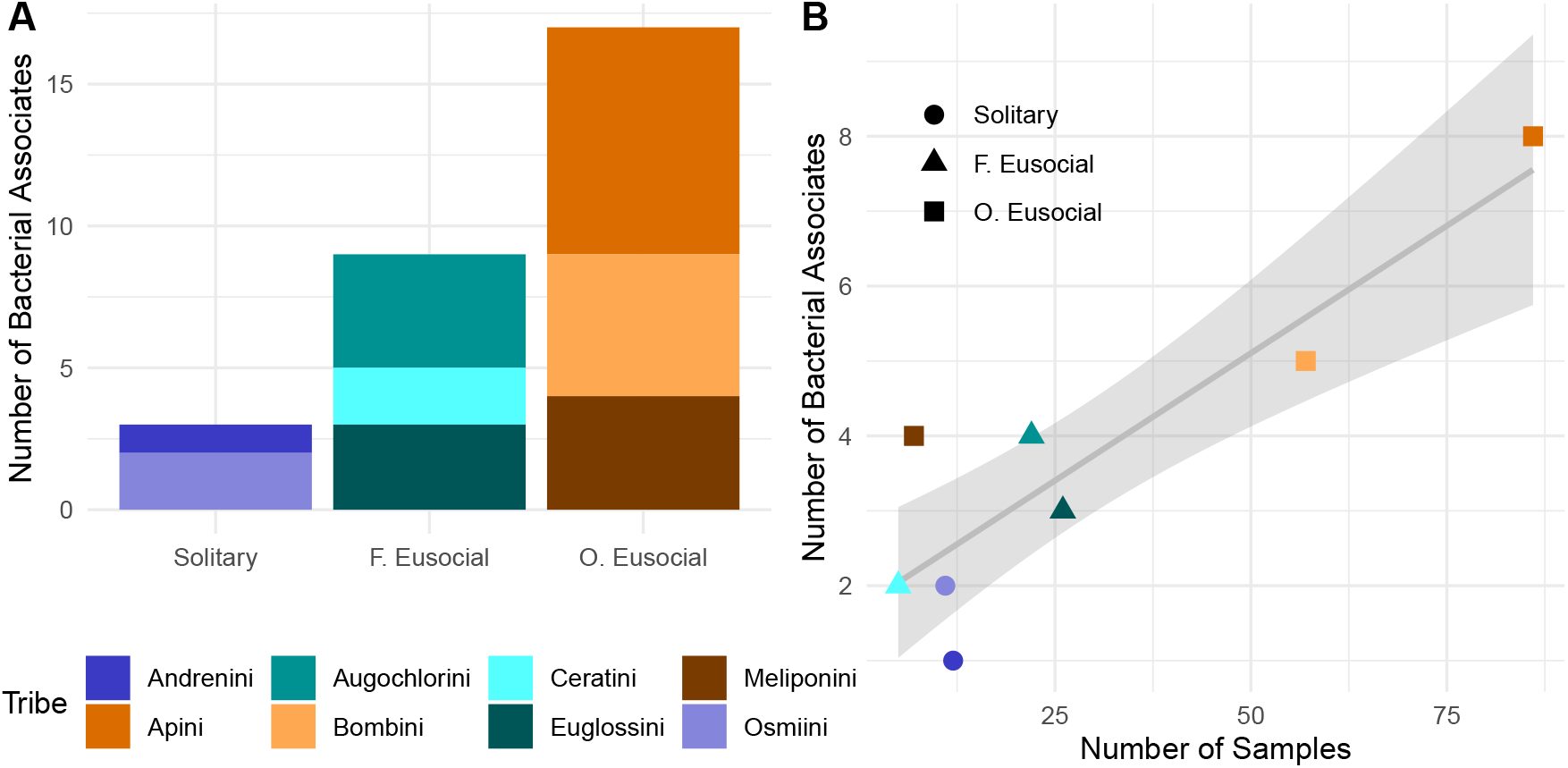
Number of bacterial associates per tribe increase with both A) sociality and B) number of samples. Meliponini has four associates from only seven samples and potentially challenges this trend. Line in plot B represents fit estimated by GLM with standard error shaded in grey.

We find *Wolbachia* associated with the two solitary tribes, Andrenini and Osmiini. *Wolbachia* was also detected at low prevalence in Apini and Bombini (Figure 2), but at comparably low average relative abundance (see **S10**). *Wolbachia* is an extremely successful insect endosymbiont, estimated to be present in as much as 52% of all insect species [Weinert et al., 2015]. This endosymbiont is capable of manipulating the reproduction of its host in order to spread throughout populations, most famously by inducing cytoplasmic incompatibility [Bourtzis et al., 1996; Werren et al., 2008], and has been proposed to be a potential factor behind *Andrena* diversification [McLaughlin et al., 2023]. In the bees, increased *Wolbachia* prevalence and diversity associated with solitary over social species has been described before [Gerth et al., 2011, 2015; Ramalho et al., 2021; Saeed and White, 2015], though the reasons for this remain speculative. As *Wolbachia* is maternally inherited, it may be that obligately eusocial societies that consist of many sterile or reproductively-constrained females would be considered an evolutionary dead-end for the symbiont, if it were not established that *Wolbachia* persists in high prevalence in a number of eusocial ant species [Ramalho et al., 2018, 2021; Russell, 2012]. It has been previously proposed that this disparity in *Wolbachia* presence between social and solitary bees occurs either due to solitary individuals having a greater number of interactions with other potentially infected taxa, or that social species have a more limited number of ecological environments within which they forage and live [Ramalho et al., 2021].

We postulate that perhaps this is more to do with obligately eusocial bees having these evolutionary long-term and stable host-microbe relationships that solitary insects are not able to achieve with their relative lack of social and inter-generational interaction. Perhaps *Wolbachia* fails to persist in social bees because the established community protects against it, at least in the case of the most social corbiculates. This phenomenon is termed “colonisation resistance” [Lawley and Walker, 2013], and many features of the social bee core microbes already identified could play a part, such as priming the host immune system [Horak et al., 2020; Kwong et al., 2017a; Lang et al., 2022; Napflin and Schmid-Hempel, 2016] or the occurrence of direct antagonistic microbe-invader interactions [Dyrhage et al., 2022; Endo and Salminen, 2013; Endo et al., 2012; Koch and Schmid-Hempel, 2012; Steele et al., 2017; Vásquez et al., 2012]. Solitary bees-such as *Andrena* species [McLaughlin et al., 2023] – missing these interconnected communities would therefore lack the protection they confer and may become vulnerable to *Wolbachia* driven reproductive manipulation. On the other hand, though often parasitic, *Wolbachia* can be advantageous to hosts conferring nutritional or fecundity benefits [Andersen et al., 2012; Cheng et al., 2019; Singh and Linksvayer, 2020] or resistance to viral or parasitic infection [Bian et al., 2010; Cogni et al., 2021; Duplouy et al., 2015; Pimentel et al., 2021; Van Den Hurk et al., 2012]. Future work testing whether *Wolbachia* are beneficial or virulent symbionts in solitary bee species would be most welcome.

### 5.3 Potential first members of shared orchid bee microbiome identified

Despite being an important group of pollinators, the orchid bees (Euglossini) remain the least studied group of corbiculate bees and, to the best of our knowledge, the microbiomes are undescribed. Two of the three orchid bee species included in this analysis – *Euglossa dilemma* and *E. viridissima* – exhibit some primitively eusocial behaviour, where a mother foundress and a subordinate daughter (sometimes two) administer brood care [Cocom Pech et al., 2008; Saleh et al., 2022]. In these instances, there is the opportunity of vertically transmitted microbes becoming established across generations, although the fact that some daughters leave the nest after eclosure would suggest these relationships could be less stable than those in obligately eusocial corbiculates. In this analysis – looking at 26 orchid bee samples – we find three Euglossini associate microbial taxa: *Asticcacaulis*, *Cupriavidus* and *Ochrobactrum* which represent the first description of the microbiota of this important group of corbiculate bees.

*Asticcacaulis* are Gram-negative bacteria that are constituents of microbial communities in freshwater, bark, and soil environments [Aschenbrenner et al., 2017; Ishizawa et al., 2019; Xie et al., 2015; Zhang et al., 2022]. Members of this genera also have potentially symbiotic relationships with plants [Jha et al., 2020; Rajkumar et al., 2009], arachnids [Hu et al., 2019; Zhu et al., 2020] and Hemipteran insects [Cooper et al., 2017]. As a plant endophyte, the genus may play a role in phosphate solubilisation, though this is yet to be definitively confirmed [Rajkumar et al., 2009]. Its role as an insect endosymbiont is not well understood and may simply be a result of its widespread presence in the environment.

*Cupriavidus* species are known for their ability to tolerate and utilize a wide range of compounds and toxic pollutants, making them important in the bioremediation of contaminated environments [Malik et al., 2021; Sohn et al., 2021]. This genus includes a well-characterised plant symbiont, *Cupriavidus taiwanensis* (previously *Ralstonia*, Chen et al. [2003]) and has otherwise been detected in the microbiome of mosquitoes [Mancini et al., 2018] and a beetle [Garcia et al., 2014]. *Cupriavidus* is a member of the Burkholderiaceae family, members of which are known to possess genomic islands that may increase its propensity for forming symbiotic relationships with insects [Stillson et al., 2022]. Despite this, and its apparent detection in other insects, *Cupriavidus* genomes assessed to date do not contain this “symbiosis” island and attempts to have other genera colonise beetle samples experimentally had uncertain results [Acevedo et al., 2021]. There is, however, evidence of these symbiosis islands transferring between different Burkholderiaceae genera via horizontal transfer [Stillson et al., 2022]. Perhaps the species we detect at genus-level here may well contain this island or similar upon more thorough sequencing. Notably, this island was detected in species of *Paraburkholderia*, which we find associated with Ceratini.

The final detected associate taxa, *Ochrobactrum*, includes genera that are known to be beneficial symbionts in plants [Babalola, 2010; Balachandar et al., 2007]. It has also been identified as an endosymbiont in snails [Dar et al., 2015], root-feeding beetles [Huang et al., 2012] and termites [Tsegaye et al., 2019; Wenzel et al., 2002]. Within these relationships, *Ochrobactrum* plays a cellulytic role, helping to break down cellulose and depolymerise lignin. *Euglossa* species enlarge tree cavities for nesting [Dressler, 1982] and perhaps to collect resin, which is an important nest building material [Cameron, 2004]. For these purposes, perhaps *Ochrobactrum*’s cellulytic abilities is potentially advantageous. This work is the first attempt at characterising the microbiome of orchid bees, and further experimental work is required to confirm these relationships and elucidate potential functions.

### 5.4 “Corbiculate” core microbes may be specific to the obligately eusocial clade

When it comes to bee microbiota, the literature overwhelmingly studies the obligately eusocial corbiculate bees, Apini, Bombini and Meliponini. This attention has identified a corbiculate core microbiome that is shared amongst these species [Kwong et al., 2017b]. These previous studies, however, do not include Euglossini, the fourth corbiculate tribe. Here we find that these core microbial taxa are not found in the same composition or prevalence as they are in the other corbiculates (Figure 4, see **S9**). Perhaps this “corbiculate” core community is a misnomer, and that what had been previously described were communities shared only between the obligately eusocial corbiculates.

There are phylogenetic implications of this insight. While the phylogeny of the corbiculates has historically been controversial, most analyses today place Euglossini as the outgroup to the other three tribes [Bossert et al., 2019; Engel and Rasmussen, 2021]. Potentially, then, this core microbiome shared between Apini, Bombini and Meliponini may be as ancient as their last common ancestor (LCA), and was composed after the split between the orchid bees and other corbiculates. It could therefore be argued that this LCA would have likely been obligately eusocial, allowing these bacterial communities to establish stably enough to be passed on to three different lineages through *∼* 55 million years of host diversification [Peters et al., 2017].

It is worth noting that our Euglossine sample size was limited (*n* = 26), and mostly consisted of *Euglossa* samples. Larger sample sizes and more species may reveal a more complicated picture of Euglossine species presenting with some or all of the “corbiculate” core microbes. However, the sample size for the Meliponini bees in this analysis was considerably smaller (*n* = 7), and yet this core community was detectable. Further microbiological investigation into the Euglossine bees would be helpful to confirm our findings.

### 5.5 Bacteria with anti-pathogen potential persist across bee taxa

Though the “corbiculate” core community was not similar between Euglossini and the classic corbiculate tribes, there were other shared microbial taxa. *Apilactobacillus* and *Bombella* / *Parasaccharibacter* – likely to actually be one genus [Smith et al., 2021] and referred to hereafter as *Bombella* – were detected in orchid bees and at considerable prevalence in *Megalopta* (Figure 4, see **S9**). Similarly, both *Apilactobacillus* and *Bombella* were detected in five and six of the ten species included in the bacterial analysis, respectively (Figure 2), though not in either of the solitary genera.

One of the reasons why these two bacterial groups are so successful at establishing in such diverse bee taxa may be their roles as anti-pathogen symbionts. *Bombella*, for example, has anti-fungal properties [Miller et al., 2021] and are found frequently in honeybee larvae and food stores, two components of the colony which are especially vulnerable to fungal infection [Anderson et al., 2014]. This would also be an advantage to any host that stores pollen, and could help explain its presence in most of the social species in this analysis. *Apilactobacillus* increases individual resistance to a number of pathogens including *Paenibacillus larvae* (American foulbrood, Butler et al. [2013]; Forsgren et al. [2010]; Kčćaniovéa et al. [2020]; Kiran et al. [2022]), the microsporidian *Nosema* [Arredondo et al., 2018], fungal infection [Iorizzo et al., 2020] and *Melissococcus plutonius* (European foulbrood, Endo and Salminen [2013]; Endo et al. [2012]; Vásquez et al. [2012]; Zendo et al. [2020]). It is also prevalent in the floral environment, suggesting an intuitive route for transmission between different bee species visiting the same flowers [Anderson et al., 2013; Tamarit et al., 2015].

Many other members of the “corbiculate” core community may also confer resistance to common bee pathogens. *Snodgrassella* increases honeybee resistance to *Serratia marcescens* infection [Horak et al., 2020], in bumble-bees *Gilliamella* and *Apibacter* suppress trypanosomatid *Crithidia* species [Cariveau et al., 2014; Mockler et al., 2018], and members of *Lactobacillus: Firm-5* inhibit *P. larvae* and *M. plutonius* growth [Killer et al., 2014] and *C. bombi* infection in bumblebees [Mockler et al., 2018]. The mechanisms of this protection could be host moderated, e.g. by increasing the expression of immune-associated genes [Horak et al., 2020; Kwong et al., 2017a], allowing for immune priming [Milutinovíc et al., 2016; Sadd and Schmid-Hempel, 2006], or symbiont moderated, e.g. by creating a physical barrier to pathogen colonisation [Kwong and Moran, 2013; Martinson et al., 2012] or producing anti-pathogen molecules [Dyrhage et al., 2022; Endo and Salminen, 2013; Endo et al., 2012; Koch and Schmid-Hempel, 2012; Steele et al., 2017; Vásquez et al., 2012].

The preponderance of anti-pathogen effects by bee associated microbes may be linked to the immune gene architecture of bees. When the honeybee genome was first sequenced [Honeybee Genome Sequencing Consortium and others, 2006], one of the curious features was the relative lack of immune genes compared to other insect models [Evans et al., 2006]. This was surprising for the honeybee, an eusocial insect that lives in societies of thousands of genetically similar individuals that are thus vulnerable to pathogen spread. Initially, this disparity was explained by the unique benefits of social immunity – a suite of behaviours that social animals use to help prevent and slow disease transmission, such as allogrooming and expulsion of the sick [Cremer et al., 2007, 2018; Dolezal and Toth, 2014; Wilson-Rich et al., 2009] – leading to relaxed selection on individual immunity and, eventually, gene loss. However, as more bee genomes became available, it became clear that this depauperate immune gene repertoire predated bee sociality [Barribeau et al., 2015].

This restricted immune genetic architecture could perhaps be why *Apilac-tobacillus* is often found outside of the classic corbiculate bees, as is found in this analysis and elsewhere [Handy et al., 2022]. In *Apilactobacillus kun-keei*, a plasmid causes one strain’s antibacterial effects against *M. plutonius* [Endo and Salminen, 2013; Zendo et al., 2020]. Upon further investigation, more plasmids putatively encoding antibiotic compounds were discovered in other strains [Dyrhage et al., 2022]. Similarly, *Apilactobacillus kunkeei* is usually found as multiple strains within hosts where transfer of mobile genetic elements are common [Tamarit et al., 2015]. These features allow for the rapid evolution of *Apilactobacillus* and may represent an example of an extended immune phenotype where the genetic potential of *Apilactobacillus* – and, perhaps, many other strains of bee-associated taxa – compensates for the relatively restricted host immune genetic potential. It is also possible that similar extended immunity phenotypes are occurring in the solitary bees – for example, the putative antiviral capability of *Wolbachia* – but these would require further investigation. It is likely that the relative lack of social-contact driven vertical transmission within the solitary species means that such relationships, when they occur, may be much more specific to solitary taxa and less permanent than what has been found in more social bees. Perhaps there are other species that, like *Wolbachia*, have evolved mechanisms to ensure high-fidelity vertical transmission without the need for consistent social interactions.

### 5.6 Mining RNA-Seq samples recapitulates experimental findings in obligately eusocial corbiculates

The composition of the “corbiculate” core microbiome has been well characterised [Engel and Moran, 2013; Engel et al., 2012; Koch and Schmid-Hempel, 2011; Koch et al., 2013; Kwong and Moran, 2016; Kwong et al., 2017b; Lim et al., 2015; Moran et al., 2012], making it a good yardstick against which we could assess the efficacy of using this pipeline to detect microbial communities. Out of the 14 microbes we opted to include as members of this core set, 12 were detected – the supposedly *Bombus*-specific *Bombiscardovia* and *Candidatus Schmidhempelia* were not detected in any samples after filtering. Having several samples per host taxa obviously improves the reliability of any detected compositions or associations, though it should be reiterated that the core microbes were recapitulated in Meliponini samples despite the relative lack of individual samples (Figure 4). We also detected the disparity in *Wolbachia* presence and abundance between social and solitary bees (Table 2), as previously described [Gerth et al., 2011, 2015; Ramalho et al., 2021; Saeed and White, 2015]. These confirmatory results were possible despite the majority of samples using poly-A enrichment as part of their library preparation (see **S1**) which reduces the level of non-eukaryotic RNA in the sample [Cui et al., 2010]. While this does limit our ability to comment on any differences in absolute abundances, this approach has consistently found the key bacterial taxa that are expected and thus represents a useful tool to estimate community composition.

While useful, this approach did not reach 100% detection of predicted microbes, and, thus, some individual microbes are potentially being missed. Despite this limitation, we was still able to unveil a number of interesting avenues for further research based on existing sequencing data. These results reveal a series of future questions that would be exciting to explore. For example, do Euglossini species lack the classic “corbiculate” core shared amongst its relatives (Figure 4)? Does it instead have specialised associations with microbes not detected elsewhere (Table 2)? Would the pattern of increasing numbers of bacterial associates with increasing social complexity hold when more solitary species are included (Figure 6)? What are the phylogenetic relationships of bacterial species with many hosts such as *Apilactobacillus*? Do obligately eusocial hosts with long-standing microbial relationships act as evolutionary reservoirs for bee symbionts, and how important are flowers in the epidemiology of gut microbial communities?

### 5.7 Conclusion

By leveraging existing RNA sequencing datasets, we was able to test whether microbial communities are affected by social structure, geography, or phylogeny. We found that bacterial community composition is significantly affected by the social lifestyle, collection location and phylogeny of the host (Figure 3, 5, see **S7-8**). In the eukaryotic and viral analyses, however, we failed to detect any factor contributing to community composition that wasn’t affected by heterogeneous dispersion (see **S7**). It appears that as the complexity of social lifestyle increases, so too does the number of bacterial associates (Table 2, Figure 6). This may be expected as prolonged social contact between host generations allows for more reliable vertical transmission and coevolution of host and symbiont. We also provide, for the first time, an initial description of the microbial community of the Euglossine bees, a complement that does not align with the regimented core microbes of their sister corbiculates (Figure 4). The anti-pathogen potential of microbial symbionts is massive, which may be how bees compensate for their own restricted immune gene arsenal. This work has highlighted many avenues that represent promising lines of future research and the need to further investigate bees of varying social lifestyles outside the classic corbiculates. Hopefully further work into the complicated, genetically mobile world of bee symbionts will further illuminate host-microbe complexities and their role in optimising bee health.

## Author Contributions

SMB and LM initially conceived the experimental approach, with SMB in particular suggesting the use of CZID.org. LM collated the samples, wrote the scripts and pipelines involved in the processing of data and underwent all analyses. LM also produced all figures used throughout this report. LM wrote the majority of the report with lots of help, guidance, patience and editing from SMB.

## Acknowledgements

Many thanks to Greg Hurst who contributed to the thinking behind this approach and Simon Hunter-Barnett who kindly imparted some of his microbiome wisdom with the authors during the analysis.

## Data Availability

The complete bioinformatic and analysis pipeline with accompanying scripts and directions are available from https://github.com/LMee17/AnthoMicroComp.

