## Supplementary Figures for "The influence of social lifestyles on host-microbe symbioses in the bees"

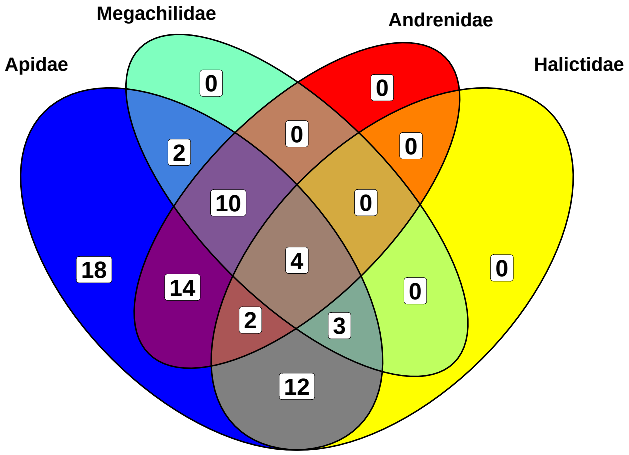


Figure 1: Overlap of bacterial taxa detected in different host families. The only unique taxa are found in Apidae.

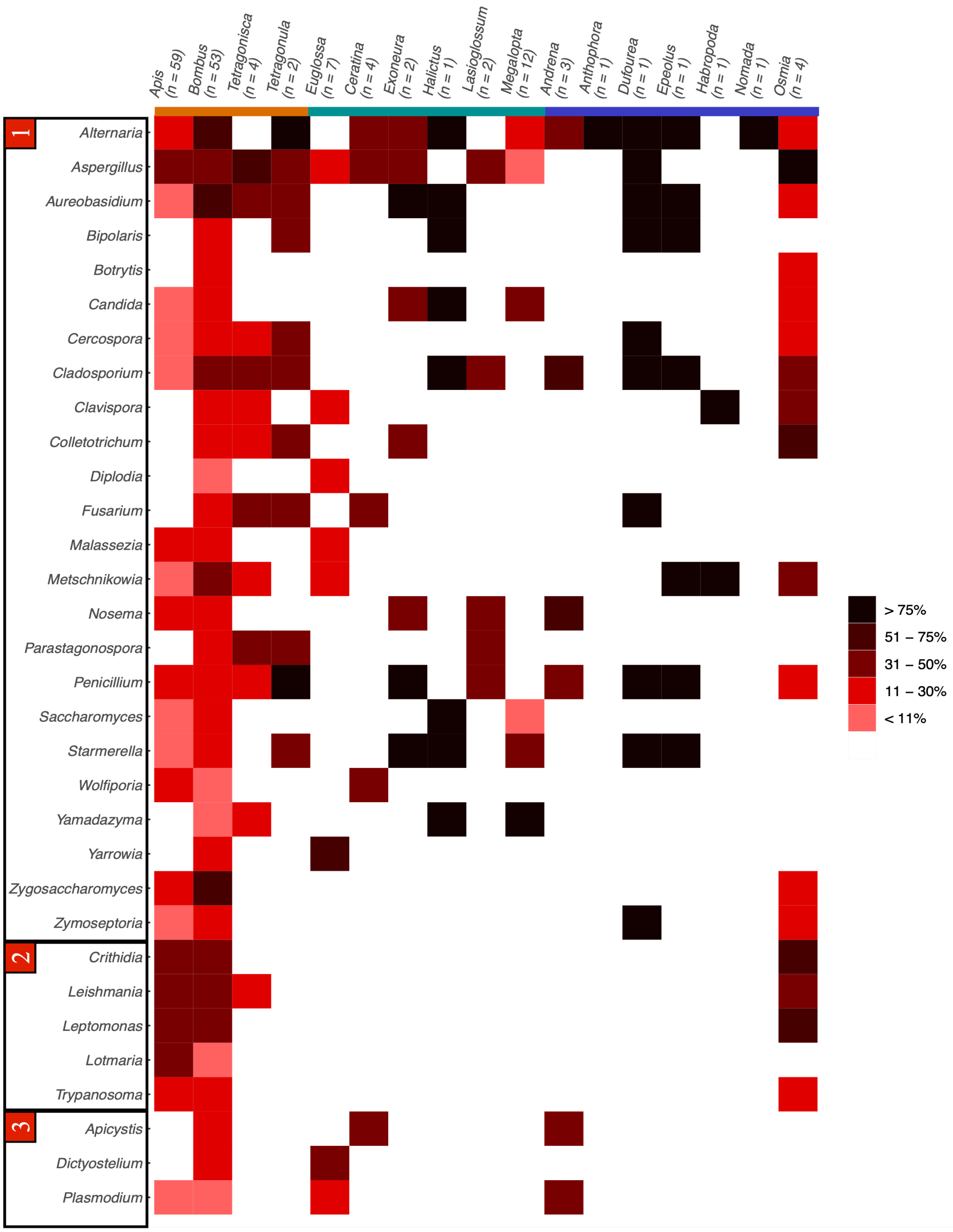


Figure 2: Heatmap of all detected eukaryote taxa and their prevalence in each genus of host samples tested after filtering. Eukaryotic taxa are ordered into 1) fungi, 2) trypanosomatids and 3) other.

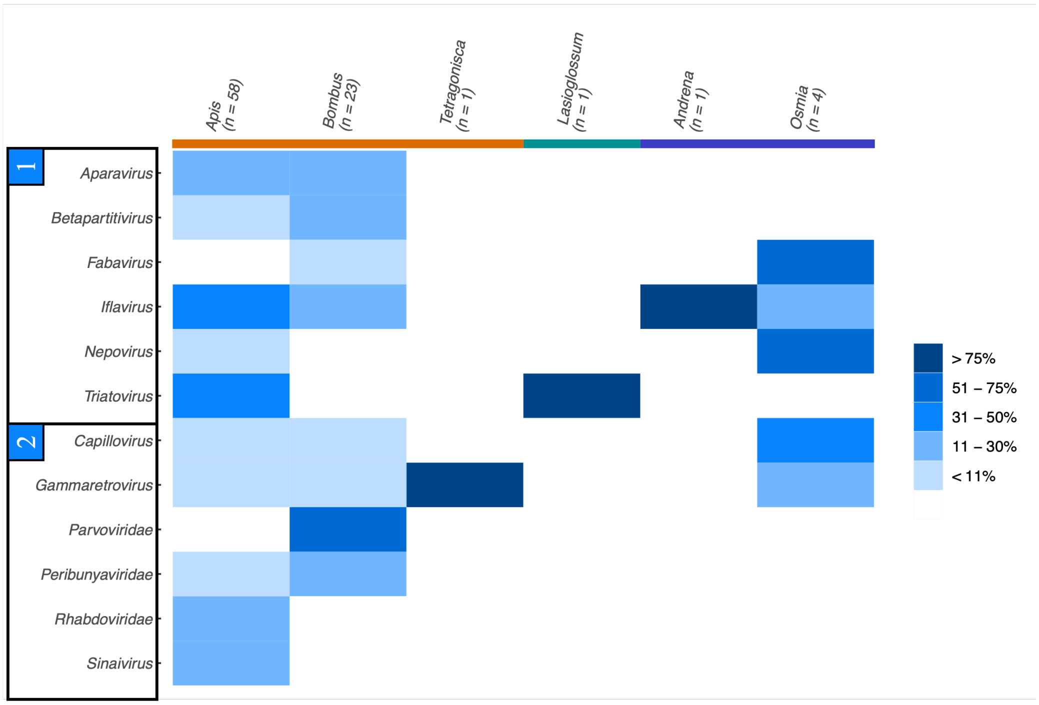


Figure 3: Detected viral prevalence in each host genus that passed data filtering. Host genera are coloured according to social lifestyle category: orange = obligately eusocial, facultatively eusocial = green, blue = solitary.
